# Gene networks under circadian control exhibit diurnal organization in primate organs

**DOI:** 10.1101/2021.09.17.460870

**Authors:** Jie Li, Pengxing Nie, Christoph W. Turck, Guang-Zhong Wang

**Author notes:** These authors contributed equally to this work (Jie Li, Pengxing Nie). Corresponding Author: Guang-Zhong Wang, Ph.D. (G-Z. Wang), Shanghai Institute of Nutrition and Health, Chinese Academy of Sciences, 320 Yueyang Road, Shanghai, China 200031.

## Abstract

Mammalian organs are individually controlled by autonomous circadian clocks. At the molecular level, this process is defined by the cyclical co-expression of both core transcription factors and off-target genes across time. While interactions between these molecular clocks are likely necessary for proper homeostasis, these features remain undefined. Here, we utilize integrative analysis of a baboon diurnal transcriptome atlas to characterize the properties of gene networks under circadian control. We found that 53.4% (8,120) of baboon genes are rhythmically expressed body-wide. In addition, >30% of gene-gene interactions exhibit periodic co-expression patterns, with core circadian genes more cyclically co-expressed than others. Moreover, two basic network modes were observed at the systems level: daytime and nighttime mode. Daytime networks were enriched for genes involved in metabolism, while nighttime networks were enriched for genes associated with growth and cellular signaling. A substantial number of diseases only form significant disease modules at either daytime or nighttime. In addition, we found that 216 of 313 genes encoding products that interact with SARS-CoV-2 are rhythmically expressed throughout the body. Importantly, more than 80% of SARS-CoV-2 related genes enriched modules are rhythmically expressed, and have significant network proximities with circadian regulators. Our data suggest that synchronization amongst circadian gene networks is necessary for proper homeostatic functions and circadian regulators have close interactions with SARS-CoV-2 infection.

## Introduction

The mammalian circadian system is hierarchical in structure, with the brain’s suprachiasmatic nucleus (SCN) acting as a master pacemaker to orchestrate clocks in other organs (Hastings et al. 2003; Welsh et al. 2010; Albrecht 2012). This system is replicated in individual cells, which need to be synchronized with each other to properly perform their organ’s functions (Dibner et al. 2010; Albrecht 2012; Mohawk et al. 2012). The central clock resets peripheral clocks slightly each day to adapt to the 24 hour day-night cycle, as the intrinsic circadian period is longer than a day (Czeisler et al. 1999). Although it is well known that a substantial fraction (3-16%) of transcribed mRNAs show rhythmic expression (Panda et al. 2002; Storch et al. 2002; Hughes et al. 2009; Zhang et al. 2014), the effects of global circadian synchronization on the transcriptomes of individual organs remains elusive.

To help understand the influence of synchronization between subordinate oscillators, we propose the concepts of “global cycling genes” and “rhythmic interactions” for analysis of the transcriptome. Global cycling genes are those transcripts which exhibit a rhythmic expression profile across multiple organs/tissues. Rhythmic interactions refer to gene pairs that are rhythmic across time. Testing these concepts has been difficult so far because only one or a few tissues are analyzed by high-throughput sequencing technologies in any given study (Doherty and Kay 2010; Hughes et al. 2017; Sun et al. 2020). Still, several large-scale circadian transcriptomes have been recently reported (Zhang et al. 2014; Mure et al. 2018; Ruben et al. 2018), making examination of this conceptual framework feasible. Through quantification of the circadian gene expression of 12 mouse organs, ~43% of protein coding genes and a large number of non-coding genes show oscillations in their expression in at least one organ (Zhang et al. 2014). Likewise, 64 baboon tissues were collected every 2 hours and utilized in RNA-seq and the data was then constructed into a high-resolution atlas of the circadian transcriptome (Mure et al. 2018). Use of such comprehensive datasets could allow examination of the molecular design principles within the circadian clock from a systemic perspective (Hogenesch and Ueda 2011).

Here, by integrative analysis of the high-resolution baboon diurnal transcriptome, we identified a large number of global cycling genes and cycling interactions, which implicate the fundamental circadian network organization of the primate body. We then characterized the co-expression properties of networks across different time points and discovered that networks can exhibit either “daytime” or “nighttime” status. Finally, we found that SARS-CoV-2 related proteins tend to be encoded by global cycling genes whose module is linked to circadian rhythms in the protein-protein interaction network. Together, these results demonstrate the effect of circadian rhythm on the whole-body primate transcriptome.

## Results

### More than half of all baboon genes are globally expressed in a rhythmic pattern

We identified global cycling genes by assessing the circadian transcriptome of more than 60 baboon (*Papio anubis hamadryas*) organs (Mure et al. 2018). No obvious outliers for any organs or circadian time points were detected based on PCA of these data (Supplemental Fig. S1). Thus, no additional normalization on FPKM value was performed to improve the detection of global cycling genes. For the 15,219 expressed genes considered, 53.4% (8,120) exhibited a significant circadian expression signal after a multiple comparison correction test (Benjamini-Hochberg adjusted *P* < 0.05) (Supplemental Table S1), suggesting that body-wide gene expression is of a robust, rhythmic nature (Fig. 1A). By removing one organ at a time, we found that >96% global cycling genes can be reproduced in more than half of the experiments (33 of 63 experiments) and >91% global cycling genes can be detected every time, which demonstrates the robustness of our methodology (Supplemental Fig. S2). By using mouse circadian atlas (Zhang et al. 2014), we conducted global cycling gene identification analysis on mice as well. We found that 50.5% of the genes (6,613 genes) in mice can be detected as global cycling genes, with a significant overlap with baboon (3,954) (Supplemental Fig. S3A, OR = 1.58, p = 1.32 x 10^-44^). With the exception of *Rora* and *Rorb*, the majority of the core circadian clock genes exhibited a significant global oscillation signal (*CLOCK*, *BMAL1*, *NPAS2*, *PER1*, *PER2*, *PER3*, *CRY1*, *CRY2*, *RORC*, *NR1D1*, *NR1D2* and *DBP*, Supplemental Fig. S4). *CLOCK* showed a weak but significant rhythmic expression profile (Benjamini-Hochberg adjusted *P* = 0.022) at body-wide level, while all other 11 core circadian genes exhibited a strong and significant oscillation signal (Benjamini-Hochberg adjusted *P* < 10^-6^) (Supplemental Table S1).

**Figure 1.**
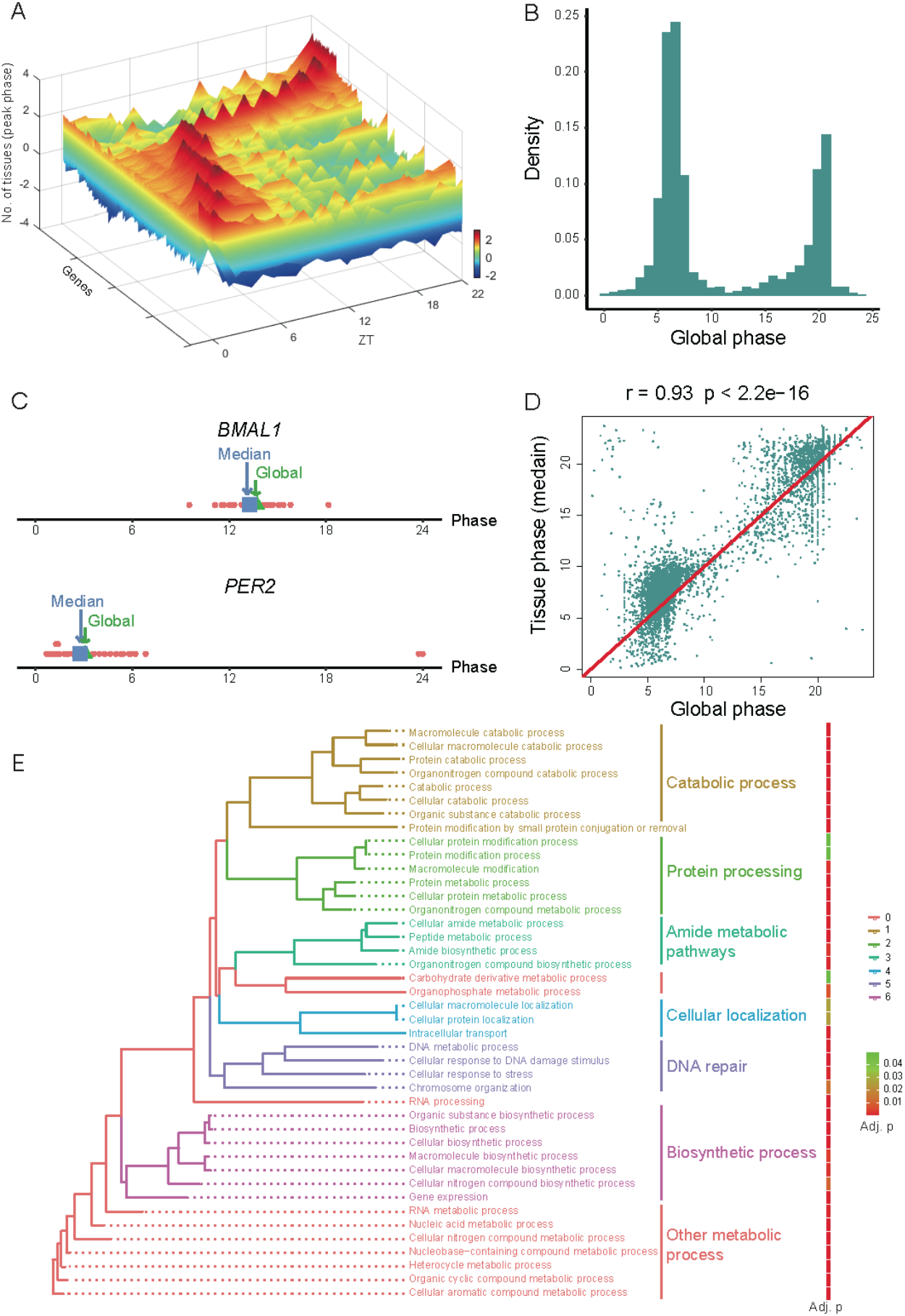
Global cycling genes and their functions. (A) 3D plot of called global cycling genes and the number of organs with peak expression. Scale bar indicates the number of organs with peak expression, with a greater number indicated in red and a lower number indicated in blue. (B) Phase distribution of global cycling genes. (C) Global phase (blue) of *BMAL1* and *PER2* are close to the median phase (green) of individual organs. (D) The correlation between global phase and median phase in individual organs for all global cycling genes. (E) The enrichment of biological processes for global cycling genes.

Consistent with previous work (Mure et al. 2018), two peak phases were observed for global cycling genes, with the first peak around noon and the second at midnight (ZT06 and ZT18, Fig. 1B). Additionally, we found that the phase of the global cycling genes represents the median phase of cycling genes in each organ (Fig. 1C, D), and thus there is a strong positive correlation between these two parameters (Pearson correlation coefficient r = 0.93, *P* < 2.2 x 10^-16^) (Fig. 1D). Subsequent functional analysis of these global cycling genes revealed that they are largely involved in basic cellular functions such as metabolic process, RNA biosynthesis, cellular localization, intracellular transport, DNA repair and others (Fig. 1E), a finding that is consistent with analysis of individual organs (Mure et al. 2018). Similar results were observed for the mouse (Supplemental Fig. S3B and C). These results indicate that the identification of global cycling genes is a useful concept that helps delineate circadian synchronization between the transcriptomes of peripheral oscillators.

### One third of gene-gene interactions exhibit rhythmic co-expression

Next, we examined to what extent gene-gene interactions exhibit rhythmic patterns. For this purpose, 10,000 gene pairs were randomly sampled from all protein-coding genes. For every circadian time point, Spearman correlation coefficients were estimated by integrating organ-based expression profiles (Fig. 2A). Thus, “gene-gene interaction” specifically indicates an association between the expression of gene pairs among diverse organs and high coefficients suggest similar expression profiles across organs. The significance of rhythmic co-expression for these gene pairs was determined by JTK_CYCLE (Hughes et al. 2010; Wu et al. 2016) and the procedure was repeated 100 times in order to obtain an unbiased assessment. 3,180 gene pairs displayed a strong signal of rhythmic co-expression (Benjamini-Hochberg adjusted *P* < 0.05) (Fig. 2B). The phase distribution of these rhythmic interactions was similar with that of global cycling genes (Fig. 2C). As the variation of individual phase increases, both the proportion of global cycling genes and rhythmic interactions decrease rather than increase (Supplemental Fig. S5A and 5B), which suggests that the rhythmicity in rhythmic co-expression cannot be explained by the variation in the cycling phase of individual tissues. For a comparison with other types of networks, a similar approach was also applied to protein-protein interactions, genetic interactions, and functional relevant interactions (genes located in the same KEGG pathway) (Kanehisa and Goto 2000; Alanis-Lobato et al. 2017; Oughtred et al. 2019; Szklarczyk et al. 2019). These data illustrated that approximately 20% of the interactions in these network types are rhythmically co-expressed, which is notably lower than that observed for the co-expression network (*P* < 2.2 x 10^-16^) (Fig. 2D).

**Figure 2.**
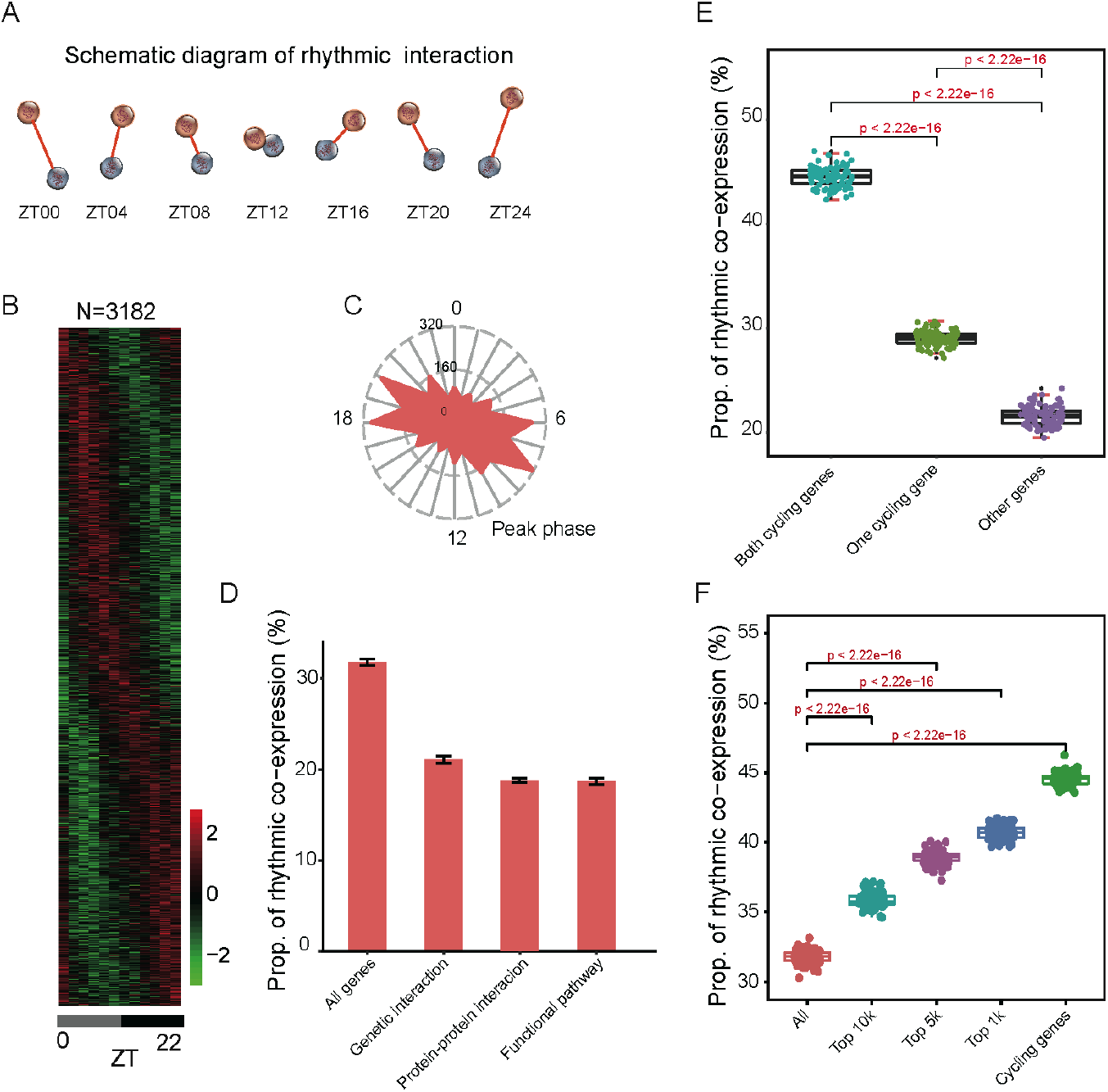
Properties of rhythmic co-expression interactions. (A) Schematic diagram of rhythmic co-expression interactions. Spearman correlation coefficients were used to measure co-expression relationships between gene pairs at given time points. Greater coefficients indicate stronger interactions between gene pairs. (B) Heatmaps of 3,182 rhythmic co-expression interactions from 10,000 randomly sampled gene pairs. Stronger interactions are marked in red while weaker interactions are marked in green. (C) Radial plot of the peak phase distribution of rhythmic co-expression links. (D) Proportion of rhythmic co-expression interactions in different networks, including protein-protein interactions (STRING), genetic interactions (HIPPIE), and functional relevant interactions (KEGG). (E) Relationships between rhythmic co-expression links and global cycling genes. Gene pairs containing both global cycling genes show the highest percentage of rhythmic interactions while gene pairs containing no global cycling genes show the lowest percentage of rhythmic interactions. (F) Relationships between rhythmic co-expression links and expression levels. Genes were ranked from highest to lowest, according to their expression levels and top 1,000, 5000, and 10,000 were grouped in the classification.

To investigate the contribution of global cycling genes on rhythmic interactions, three situations were considered: (i) when both are global cycling genes, (ii) when only one is a global cycling gene, and (iii) when neither one is global cycling gene. For each category we examined the proportion of rhythmic interaction. For 10,000 randomly selected gene pairs in case (i), 4,450 interactions were rhythmically co-expressed, while in cases (ii) and (iii) only 2,903 and 2,158 gene pairs exhibited rhythmic links, respectively (*P* < 2.2 x 10^-16^) (Fig. 2E). As cycling genes are highly expressed (Wang et al. 2015; Cheng et al. 2019), we then sampled the same number of gene pairs from the top 1,000, 5,000 and 10,000 expressed genes. We found that a considerably greater percentage of rhythmic co-expression links were detected in these groups than expected by chance (*P* < 2.2 x 10^-16^) (Fig. 2F). Such results demonstrate that global cycling genes substantially contribute to bodywide rhythmic interactions.

Circadian oscillation is generated by the transcriptional/translational autoregulatory feedback loop (TTFL). The core circadian network consists of *CLOCK* (along with its paralog *NPAS2*) and its heterodimeric partner *BMAL1* (*ARNTL*), which bind to their downstream targets including *PER1*, *PER2*, *CRY1*, and *CRY2* (Partch et al. 2014; Takahashi 2017; Rijo-Ferreira and Takahashi 2019). We assessed whether the interactions of the members of the core circadian regulatory network exhibit a rhythmic co-expression pattern. Strikingly, *CLOCK* and *BMAL1* are robustly and rhythmically co-expressed, with a peak phase at ZT13 (Benjamini-Hochberg adjusted *P* = 4.22 x 10^-5^), indicating that the two core circadian partners bind to each other in a time dependent manner. For the potential interactions amongst the 14 core circadian genes, 62.22% (56/91) were observed to exhibit a robust rhythmic interaction, which was significantly higher than the proportion of other detected interaction pairs (Supplemental Table S2; Supplemental Fig. S6). The high proportion of rhythmic co-expression among core circadian genes implies that the periodicity of these interactions may be a fundamental feature of core circadian oscillators (Wallach et al. 2013).

### Rhythmic expression of body-wide network modules

Because the co-expression of gene pairs results in distinct gene clusters that form network modules (Hartwell et al. 1999; Barabasi and Oltvai 2004), we asked whether the latter also exhibit rhythmic characteristics. Weighted Gene Co-expression Network Analysis (WGCNA) was constructed by using all the samples from the circadian expression atlas (Langfelder and Horvath 2007; Langfelder and Horvath 2008), with 20 modules detected (Fig. 3A, B; Supplemental Table S3). 9 modules exhibited significant rhythmic signals based on either their module eigengenes or the average expression of each module (brown, yellow, cyan, tan, midnight blue, black, red, turquoise, salmon modules, Benjamini-Hochberg adjusted *P* < 0.05) (Fig. 3C; Supplemental Table S3). The peak phases of 5 of these rhythmic modules were around noon (brown, yellow, cyan, midnight blue, black modules), while the peak phases of other 4 rhythmic modules occurred around midnight (tan, red, turquoise, salmon modules), suggesting that phase peak times also impact network modules. We also found that 7 of the 9 rhythmic modules are enriched for global cycling genes (Fig. 3D). Additionally, 3 core circadian genes (*PER1*, *NR1D1*, and *DBP*) were found in the “brown” cycling module, whereas 6 core circadian genes (*NR1D2*, *CLOCK*, *RORA*, *PER3*, *RORC*, and *BMAL1*) were located in the “turquoise” cycling module. These results indicate that circadian oscillation is a basic property of body-wide gene networks.

**Figure 3.**
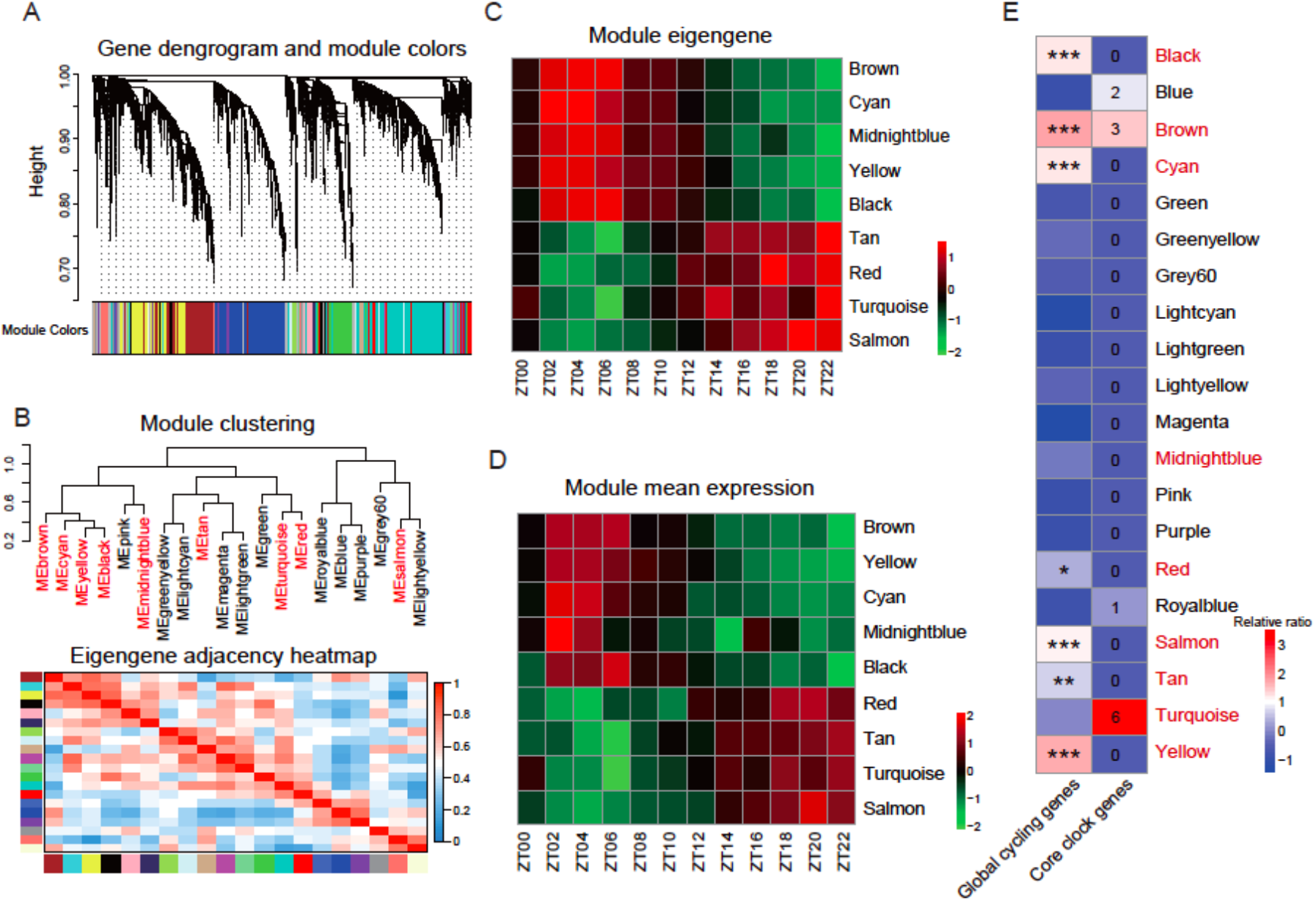
Exhibition of network rhythmicity at modular level. (A) Gene cluster dendrograms and module dictation using all expressed genes in the circadian transcriptome atlas. (B) Similarity between network modules. Rhythmic modules are indicated in red. (C and D) Heatmap of the eight rhythmic modules indicated by eigengene or mean expression. (F) The relationship between rhythmic modules and global cycling genes. Blocks with stars indicate the significance of the enrichment (* *P* < 0.01, ** *P* < 0.0001, *** *P*< 1e-10; Fisher’s exact test), and blocks with numbers indicate the number of core circadian genes included in each module.

### Co-expression networks are more internally connected at nighttime than during daytime

To investigate the dynamics of body-wide networks across time, we then built detailed WGCNA modules for each circadian time point and computed the network topology accordingly (Horvath and Dong 2008). In order to make different networks comparable, similar parameters were used (soft threshold = 16) for each network’s construction (Supplemental Codes). We found that these networks exhibited distinct properties (Supplemental Table S4). The median network connectivity at night is significantly higher than during daytime (Average degree = 38 vs. 35, respectively, *P* = 0.013) (Fig. 4A), suggesting that the expression profiles of different organs are more similar to each other at night. The network density fluctuated about 54% between daytime and night, and the network is denser at night than during daytime (0.00515 vs. 0.003, respectively, (*P* = 0.0076) (Fig. 4B). The differences found between nighttime- and daytime-assigned networks were preserved when using other network parameters, such as centrality (*P* = 0.0076) (Fig. 4C), cluster coefficients (*P* = 0.021) (Fig. 4D), heterogeneity (*P* = 0.021) (Fig. 4E), and Maximum Adjacency Ratio (*P* = 0.0022) (Fig. 4F). Overall, the topological differences suggested that the body-wide co-expression networks are flexible during daytime and robust at night (Fig. 4G-I).

**Figure 4.**
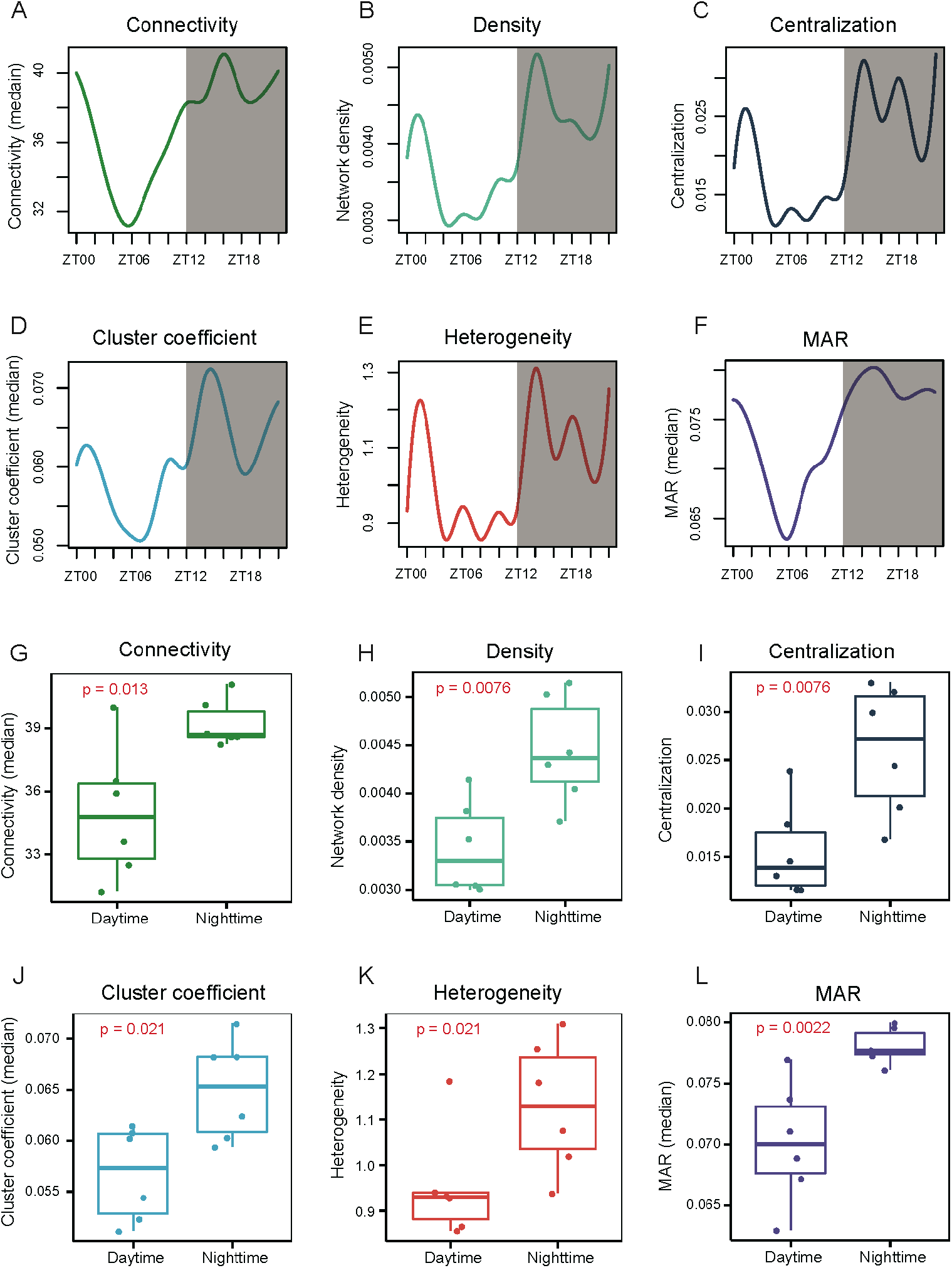
Comparison of network topology at daytime and nighttime. (A-F) Change in network density, centralization, heterogeneity, connectivity, cluster coefficient, and maximum adjacency ratio (MAR) across different time points. Daytime phase is indicated in white and nighttime phase is indicated in gray. (G-I) Comparison of network topological parameters between daytime and nighttime networks (Wilcoxon’s rank-sum test).

### Networks at the two peak phases impact distinct functions

To further characterize the differences between daytime and nighttime networks, we focused on the phase peaks at ZT06 and ZT18 (Fig. 1B), which correspond to noon and midnight, respectively. The midnight (ZT18) networks possessed both a higher connectivity and clustering coefficient than the daytime networks (ZT06) (Fig. 5A, B). 43 and 42 network modules were classified in each network, respectively (Fig. 5C, D; Supplemental Table S5). Although the two network types exhibited distinct network topologies (Fig. 5A, B), approximately half of the modules detected were shared between them (Supplemental Fig. S7). Many of these “consensus” modules were related to a specific organ (Supplemental Fig. S8; Supplemental Table S6), indicating the conservation properties of these two network types. Brain, muscle, and immune system tissues were associated with five, four, and four consensus modules, respectively (Supplemental Table S6), and included 5,223 global cycling genes in total. We next examined the differentially connected nodes of these two network types by focusing on genes with a network connectivity that is at least 2-fold higher in one network than the other. Of the 5,755 differentially connected genes, 3,767 (65%) were global cycling genes (Fig. 5E), which were over-represented (Fisher’s exact test, *P* = 2.05 x 10^-122^). In addition, we also determined the Euclidean distance for each gene between ZT06 and ZT18 by taking the co-expression weights into consideration. We found that global cycling genes have greater Euclidean distance compared to other genes, indicating that global cycling genes tended to have different network partners than other genes in the two networks (*P* < 2.2 x 10^-16^) (Fig. 5F). Subsequently, we asked whether global cycling genes can explain the distinct strength of network connections. We found that global cycling gene pairs displayed greater differences in co-expression coefficients compared with cases where only one or none of the two interacting partners was a global cycling gene (*P* < 2.2 x 10^-16^).

**Figure 5.**
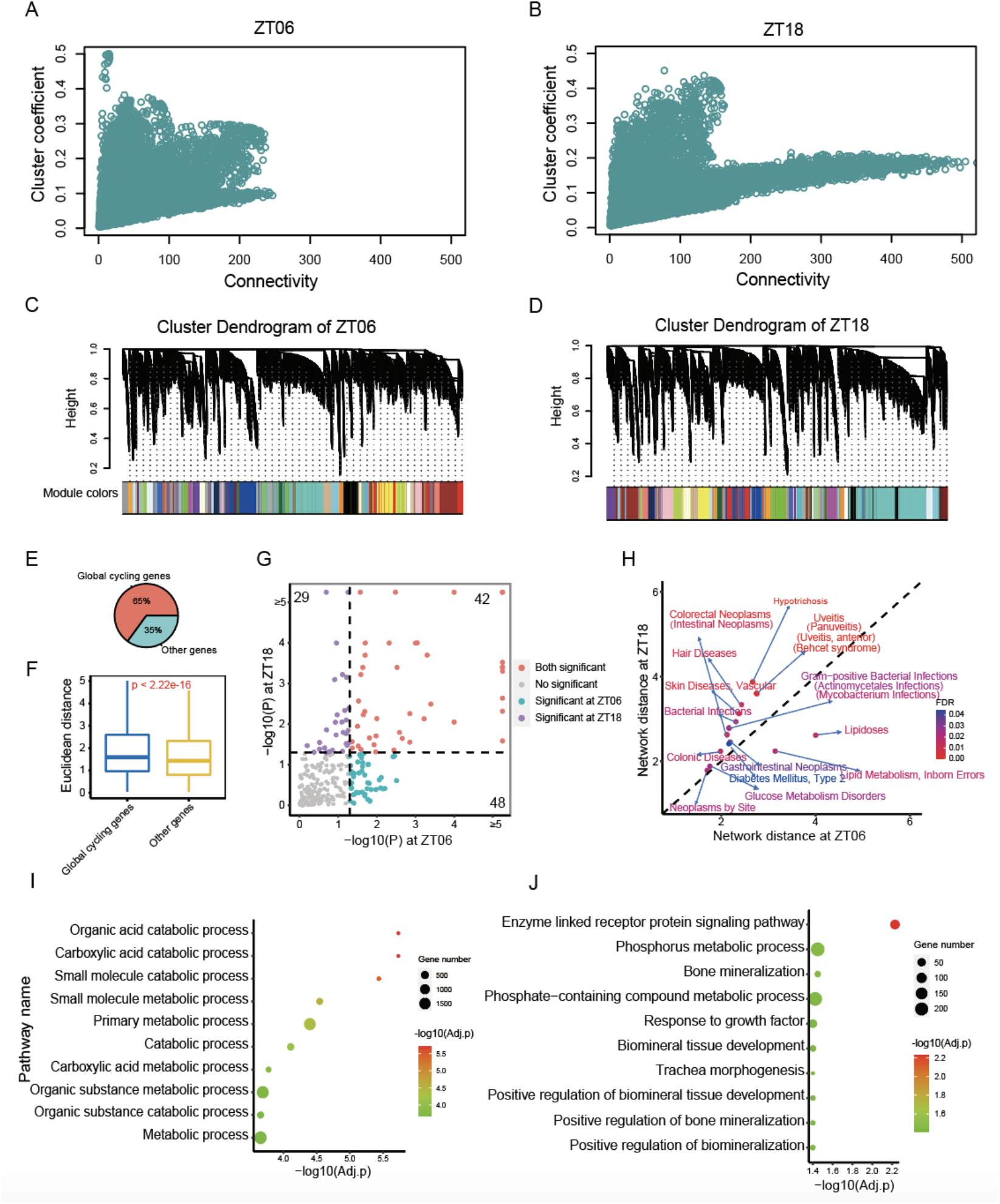
Network comparisons between phase peaks at noon (ZT06) and midnight (ZT18). (A and B) The relationship between connectivity and cluster coefficients at ZT06 and ZT18. (C and D) Gene clustering dendrogram and module distributions at ZT06 and ZT18, of which 43 and 42 modules were assigned, respectively. (E) The proportion of global cycling genes and non-cycling genes of all differentially connected genes. (F) Euclidean distance between global cycling genes and non-cycling genes within the networks. (G) Significance of the disease modules at ZT06 and ZT18. (H) The network-based distance of representative disease modules at ZT06 and ZT18 and the statistical significance of the difference. (I) Top 10 enriched biological pathways of genes with less connectivity at ZT06. (J) Top 10 enriched biological pathways for genes with less connectivity at ZT18.

Disease module is an important concept in network medicine, implicating the pathology of complex diseases and their related drug partitioning(Barabasi et al. 2011; Menche et al. 2015). The daytime and nightttime networks implicate that those disease modules are not static. Indeed, we found that many diseases only form significant disease modules at either daytime or nighttime, *i.e*. disease modules formation depends on the network dimorphism (Figure 5G). We have plotted 20 representative diseases modules according to whether they are daytime or nighttime oriented. Interestingly, it appears most of the diseases are nighttime oriented (Figure 5H), such as glucose metabolic disorders (Supplemental Fig. S9). This may have important implications for treatment and disease pathology. Finally, we found that >30% of approved drug target genes in the U.S. Food and Drug Administration Drug Bank (www.drugbank.ca) represent network hubs that are differentially expressed between daytime and nighttime networks, which further implicates the biomedical significance of the two circadian network statuses.

Besides the differences in network topology, we also found that the daytime and nighttime networks are associated with distinct functions and organs may cycle through functions throughout the day. We then focused on the genes that have less connectivity with other genes, as such these genes could indicate dissimilarities in organ activities across circadian time. Genes with fewer network partners at ZT06 (daytime) were associated with various metabolic processes (including *organic acid catabolic process*, adjusted *P* = 1.86 x 10^-06^; *carboxylic acid catabolic process*, adjusted *P* = 1.86 x 10^-06^; *small molecule metabolic process*, adjusted *P* = 2.84 x 10^-05^ and *organic substance metabolic process*, adjusted *P* = 0.0002) while genes with small amounts of network connectivity at ZT18 (nighttime) were implicated in growth-related functions (*bone mineralization*, adjusted *P* = 0.035, *response to growth factor*, adjusted *P* = 0.040 and *biomineral tissue development*, adjusted *P* = 0.040) or cellular signaling pathways (*enzyme linked receptor protein signaling pathway*, adjusted *P* = 0.0059) (Fig. 5I,J; Supplemental Fig. S10; Supplemental Table S7). Thus, the metabolic functions of specific gene modules were more dynamic during daytime, seemingly fitting the energetic demands of diurnal animals. Interestingly, five core circadian genes (*BMAL1*, *PER1*, *PER2*, *DBP*, and *RORB*), all of which cycled in a body-wide pattern, displayed at least two-times higher connectivity at night (ZT18) than at daytime (ZT06), implicating the core circadian clock in this is transcriptomic architecture.

### Global cycling genes and rhythmic interactions tend to be COVID-19 related

The COVID-19 pandemic has resulted in hundreds of millions of infections with millions of those being fatal (World 2020; Zhu et al. 2020). The relationship between SARS-CoV-2 infection and circadian regulation has not been fully explored. We thus examined the recently identified 332 human proteins that are reported to interact with SARS-CoV-2 (Gordon et al. 2020). Surprisingly, we found that 69.01% (216) of these genes were global cycling genes (Fisher’s Exact Test, OR = 1.97, *P* = 1.57 x 10^-8^). This enrichment cannot be explained by the high expression level of these genes (*P* = 3.79 x 10^-8^, logistic regression analysis). Further, among these 216 cycling genes, 92 exhibited oscillating expression in 13 human tissues (Ruben et al. 2018) and 61 of them showed periodic expression in at least one organ in both mouse and human (Zhang et al. 2014; Ruben et al. 2018) (Supplemental Table S8). Enrichment analysis using the Human Protein Atlas suggested that the 61 conserved global cycling genes were highly expressed in respiratory epithelial cells, bronchi, the placenta, the epididymis, glandular cells, the kidneys, and cells in tubules (adjusted *P* < 0.001). In addition, 52.06% of co-expression interactions among those 216 global cycling genes were highly rhythmic, which was greater than for gene pairs from the entire set of 313 SARS-CoV-2 interacting genes (42.68%) or from all the global cycling genes (44.52%) in the co-expression network.

To validate the enrichment between global cycling genes and SARS-CoV-2 related proteins, we explored an independent dataset, *i.e*. SARS-CoV-2 host factors (Daniloski et al. 2021; Schneider et al. 2021; Wang et al. 2021; Wei et al. 2021). It appears that this enrichment still exists (Fisher’s Exact Test, OR = 2.10, *P* = 2.41 x 10^-10^) and cannot be explained by expression level (*P* = 9.55 x 10^-10^, logistic regression analysis, Figure 6A). We also found that >80% of the SARS-CoV-2 related genes enriched network modules are rhythmic modules (Figure 6A), whose functions are tightly associated with autophagy (*P* = 0.0017), viral process (*P* = 0.0087) and intracellular transport (*P* = 6.43 x 10^-12^) (Figure 6B, 6C and 6D). Furthermore we found that for the differential connected SARS-CoV-2 interacting proteins between ZT18 and ZT06, > 90% of them are global cycling genes (33 out of 36 genes, 44 out of 48 genes, respectively).

**Figure 6.**
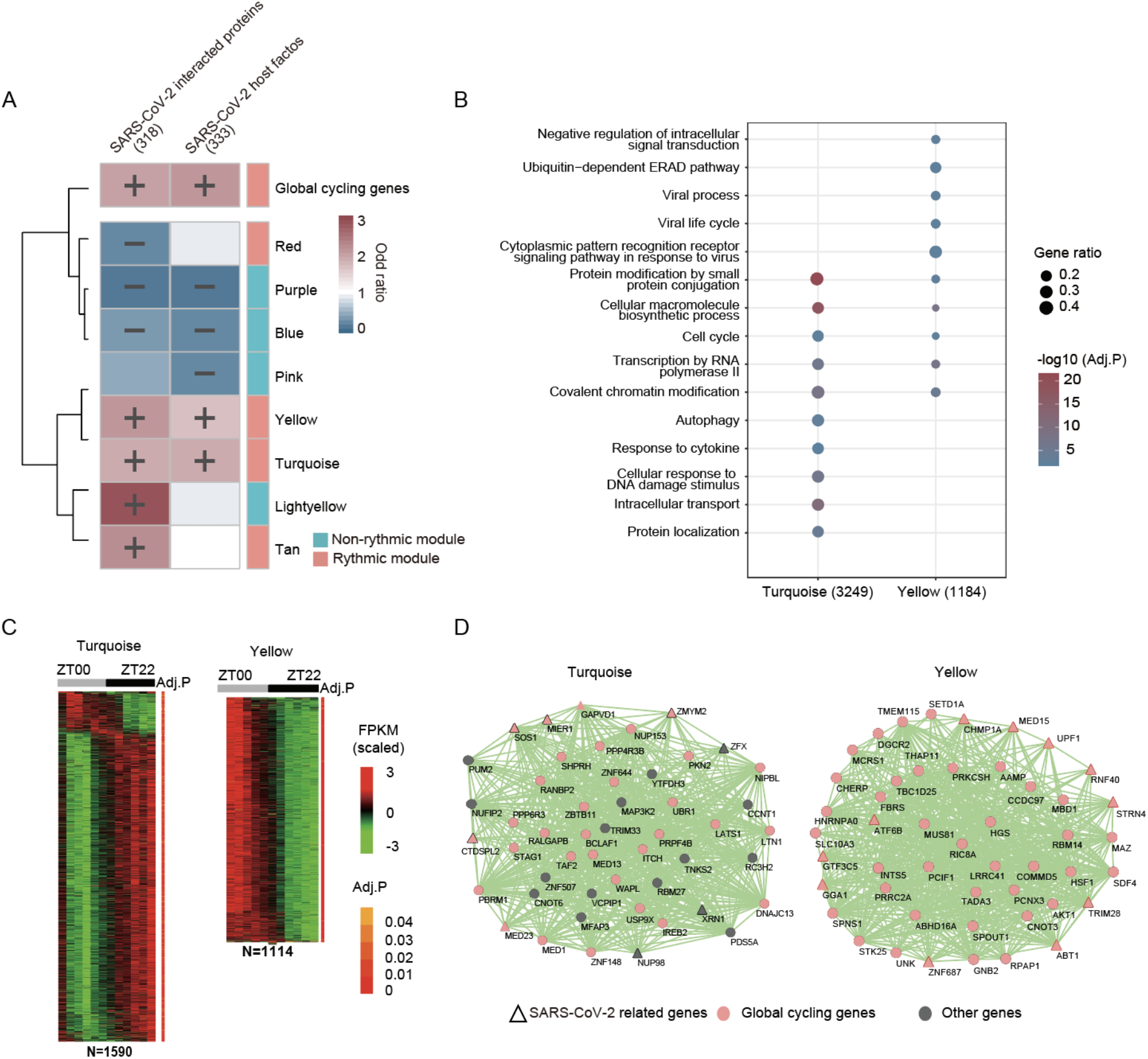
Relationship between SARS-CoV-2 related genes and diurnal modules. (A) Heatmap shows the over-representation of SARS-CoV-2 related genes in global cycling genes and diurnal modules. (B) Functional enrichment of rhythmic modules that are enriched in SARS-CoV-2 related genes. (C) Heatmap representation of the global cycling genes in module yellow and turquoise. (D) Network view of the top 50 genes defined by the highest intramodular connectivity in module yellow and turquoise. Pink nodes are global cycling genes, dark grey nodes are non-global cycling genes, while SARS-CoV-2 related genes are shown as triangles.

Most importantly, the module of circadian regulation and SARS-CoV-2 interacting genes (either host factors or SARS-CoV-2 interacting proteins) are significantly closer in the protein-protein interaction network (*P* = 0.011 and 0.020, respectively, Supplemental Table S9), suggesting that SARS-CoV-2 infection may disrupt the circadian regulatory pathway as well. Indeed, many proteins that surround the two network modules are related to viral infection process (*P* = 2.9 x 10^-119^ and 4.2 x 10^-123^, Figure 7A-C) and immune response (*P* = 0.0079 and 0.0154, Figure 7B and 7D, Supplemental Table S9). We thus posit that the treatment of circadian disruption may also be beneficial for the recovery of SARS-CoV-2 infection and the relief of the long COVID symptoms.

**Figure 7.**
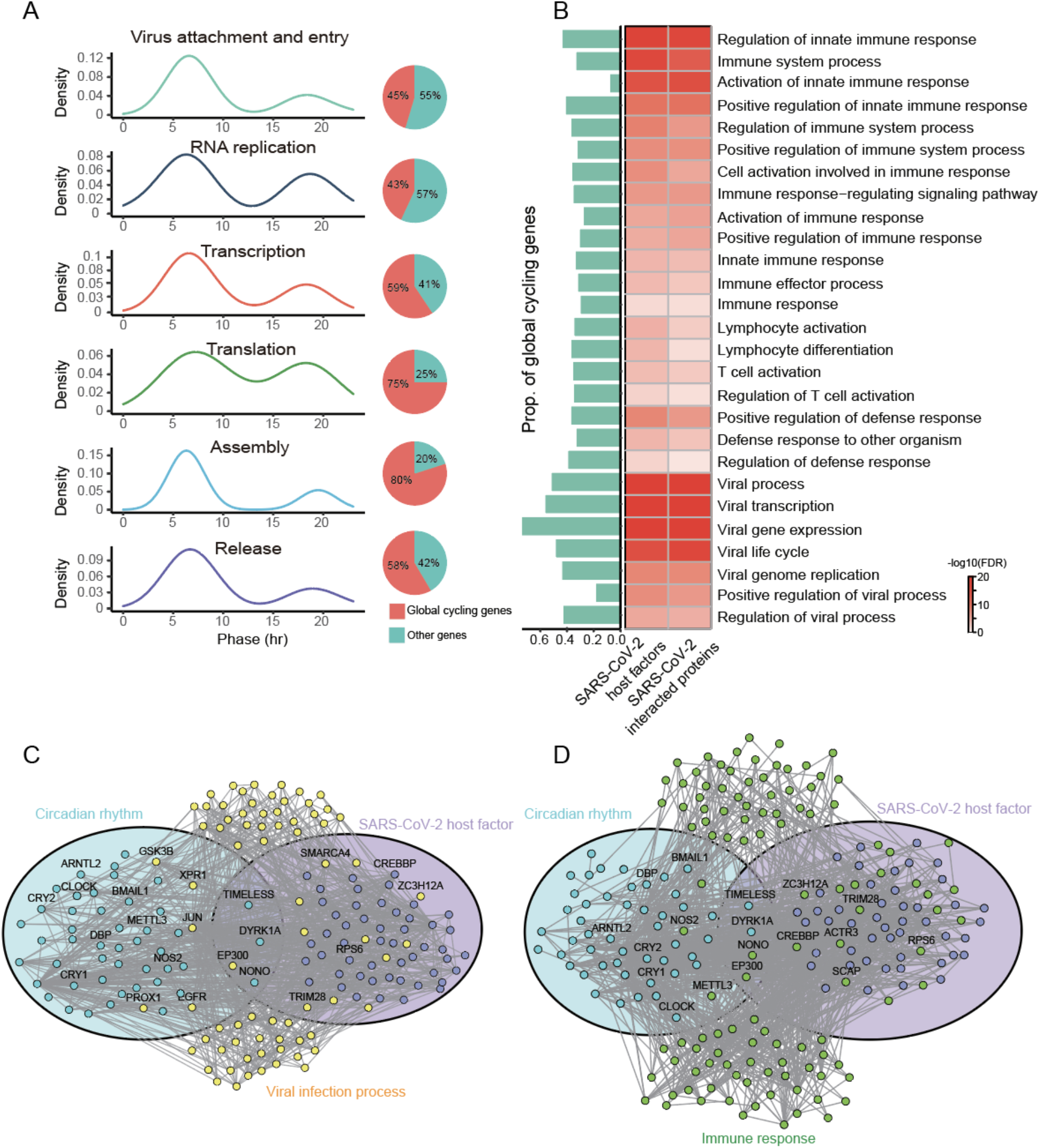
Network-based model of SARS-CoV-2 related genes and circadian regulation. (A) Phase distributions and proportion of global cycling genes involved in different processes of viral infection. (B) Functional annotation of internodes between “circadian rhythm” and “SARS-CoV-2 host factors” or “SARS-CoV-2 interacted proteins”. (C-D) A network landscape of circadian rhythm related genes and SARS-CoV-2 host factors, with genes involved in viral process (C) and immune response (D) surrounding them. Light blue circle represents the genes whose function is directly related to circadian rhythm, and genes in light purple circle are SARS-CoV-2 host factors. Yellow nodes are “viral process” related genes and green nodes are “immune response” related genes.

## Discussion

Individual organs require coordination with other organs to perform unified functions and ensure homeostasis. Similarly, individual cells in an organ need to be synchronized in order to implement the function of that organ, and this relies on gene co-expression. Hence, two genes that are related in function need to be properly co-expressed across time. By analyzing body-wide diurnal transcriptome data from >60 baboon organs, we found that more than half (53%) of transcripts are rhythmically expressed on a global scale. The phase of these genes represents the median phase of the cycling genes in each organ. More importantly, two modes of network status were discovered, with daytime networks associated with metabolic functions and nighttime networks associated with growth-related processes. At least one third of interactions and half of all network modules were rhythmic, revealing a cyclical nature to organ-specific output. Interestingly, this rhythmicity is widely enriched in genes encoding SARS-CoV-2 interacting proteins.

Several novel aspects of circadian rhythms were revealed by our approach of evaluating global cycling genes and rhythmic interactions. Firstly, by integrating the circadian transcriptomes of multiple organs, robust detection of cycling genes was achieved with high-confidence (Benjamini-Hochberg adjusted *P* < 0.05), even though the sequencing depth of samples was relatively low (median total of 18.7 million sequence reads per sample). Secondly, this approach allowed detection of coordination between multiple oscillators, resulting in dynamic regulation of thousands of genes, which has been demonstrated in both baboon and mouse. Thirdly, most of the core circadian genes including *CLOCK* exhibited oscillation at the body-wide level, which is distinct from their behavior in individual tissues (Mure et al. 2018); lastly, the interactions among human SARS-CoV-2 interacting genes were also rhythmic and have close links with circadian regulation.

The nature of gene networks has been extensively studied and several important features have been revealed. For instance, ‘small-world’ and ‘scale-free’ features characterize the structural organization of biological networks (Watts and Strogatz 1998; Barabasi and Albert 1999). Hub genes in the network frequently indicate portend the functional importance of a particular gene (Konopka et al. 2012; Yang et al. 2014). Our data show that rhythmicity is another fundamental feature of biological networks. This is property might be extended to protein-protein interaction networks, genetic interaction networks, functional association networks and regulatory networks, although the proportion of the rhythmic interactions each network could differ. The rhythmicity of biological networks reveals that gene networks may function in a temporally-organized manner, allowing different parts of the network to work sequentially.

Overall, co-expression networks were significantly more internally connected at night suggesting that there are two basic network modes: daytime and nighttime mode. This network dimorphism implicates that circadian clock is regulating metabolism to assist feeding/digestion during the day and tissue repair/growth at night and may have biomedical implications, Recent work implies that circadian time should be considered as a factor during drug development, as it may affect drug efficiency, especially for those with short half-lives (Cederroth et al. 2019; Panda 2019; Ruben et al. 2019). Indeed, we found that many diseases only form disease modules either at daytime or nighttime. Since network dimorphism indicates the similarity of drug target activities in these organs, we submit that for circadian medicine, these two network modes need to be validated and then tested for their usefulness in drug development and usage.

Although the cycling genes display two phase peaks, the time point of these phase peaks can differ between genes (Zhang et al. 2014; Mure et al. 2018). Moreover, the diurnal nature of such peaks have remained elusive. By constructing peak phase co-expression networks, we have found that daytime- and nighttime-assigned networks have differential topologies and associated functions, with greater regulation of metabolic pathways during daytime and more regulation of growth and cell-signaling functions at night. The function of these peak phases is consistent with the daily activity of baboon.

The inflammatory course of viral infections are closely linked to the circadian system (Edgar et al. 2016; Gachon et al. 2018; Hong et al. 2018; Sengupta et al. 2019), especially for influenza. The rhythmic expression of genes encoding SARS-CoV-2 interacting proteins and host factors together with their enrichment in rhythmic modules shows that SARS-CoV-2 may be associated with circadian rhythms, which has potential implications for health care efforts (Liu et al. 2020; Zhang et al. 2020). The organs with the most circadian-based regulation of SARS-CoV-2 interacting genes may be the ones where the response to SARS-CoV-2 infection is time dependent. More importantly, SARS-CoV-2 related proteins have closer network proximity with circadian rhythm than expected by chance. Many of the symptoms of long COVID(Blomberg et al. 2021), like extreme tiredness, poor concentration, difficulty sleeping, headaches and depression, are similar with symptoms observed for circadian rhythm disorders. This would imply that SARS-CoV-2 patients should increase their “circadian robustness” by avoiding behaviors that disrupt this system, such as insufficient sleep and excessive fatigue. Time restricted eating may be benefit to the recovery as well. The design of the COVID-19 vaccination may also need to take this chronobiology factor into consideration (Long et al. 2016) and thus this represents an rich area for future investigation.

In conclusion, we find that both global rhythmic expression patterns and interaction signatures are robust features of gene networks; thus reflecting circadian synchronization amongst different organs and tissues. We show that such models can be successfully applied to the analysis of single cell oscillations, as the transcriptome of individual cells should be aligned together with the cycling functions of their particular organs. Forthcoming research that applies single-cell sequencing technologies to the molecular architecture of the circadian clock could help validate our findings herein (Wen et al. 2020).

## Methods

### Primate diurnal gene expression atlas

FPKM expression values from 756 samples were downloaded from NCBI’s Gene Expression Omnibus, with GEO accession number: GSE98965 (Mure et al. 2018). For each organ, only genes with expression levels above 0 for at least half of all the time points were considered to be expressed. In the final dataset, the expression profiles of 15,219 genes were included for further analyses.

### Baboon global cycling genes identification

To detect global cycling genes, all the expression information from 63 organs was considered at a particular time point. JTK_CYCLE (Hughes et al. 2010; Wu et al. 2016) was employed to detect the global rhythmic parameters, such as amplitude, phase period, and the significance level. Only genes with a Benjamini-Hochberg adjusted *P* value < 0.05 were considered to be global cycling genes.

### Mouse global cycling gene identification

We downloaded the circadian microarray data of 12 mouse tissues from NCBI (GSE54650) (Zhang et al. 2014). Then the expression levels of 13,087 mouse genes that are homologous with baboon genes were extracted. Next, ComBat was used to eliminate the variations among organs (Johnson et al. 2007) and the JTK_CYCLE was used to detect global cycling genes. Only genes with Benjamini-Hochberg adjusted *P* < 0.05 were considered to be global cycling genes.

### Rhythmic interaction identification

10,000 gene pairs were randomly sampled from the expressed genes. For each gene pair, the Spearman correlation coefficient was estimated based on expression among 63 organs. Then, JTK_CYCLE was used to estimate the rhythmic parameters and only gene pairs with Benjamini-Hochberg adjusted *P* values < 0.05 were retained as rhythmic interactions. The above procedures were repeated 100 times to obtain an unbiased assessment. Protein-protein interactions were obtained from the STRING (Szklarczyk et al. 2019) and HIPPIE (Alanis-Lobato et al. 2017) databases; genetic interactions were downloaded from the BioGRID (Oughtred et al. 2019) database, and functional relevant interactions were downloaded from the KEGG pathway database (Kanehisa and Goto 2000). Then the proportion of rhythmic co-expression in these types of interactions was analyzed via JTK_CYCLE.

### Co-expression network analysis

To investigate rhythmic co-expression characteristics at the module level, co-expression networks were constructed based on the expression data among diverse organs. In brief, Pearson correlation coefficients were used to calculate expression similarities across tissues. Then, the power parameter was selected as 16. In total, 15,219 genes were assigned into 20 modules. To estimate whether one particular module is rhythmic, both the module eigengene and the mean expression of this module were analyzed by JTK_CYCLE. Modules with Benjamini-Hochberg adjusted *P* values < 0.05 were considered to be rhythmically expressed.

Then we applied the same soft threshold (16) to build the co-expression network at each time point. To compare the network topologies at different time points, six network indices were calculated by the building function *fundamentalNetworkConcepts* in WGCNA (Langfelder and Horvath 2008); including network connectivity, cluster coefficient, maximum adjacency ratio (MAR), network density, centralization and heterogeneity.

### Co-expression network comparison at noon and midnight

To compare the peak phase networks, which occur at noon and midnight, the above expression networks built at ZT06 and ZT18 were chosen for downstream analyses, respectively. Differentially connected genes were defined as genes with > 2-fold changes in their connection degree in the two networks. Then the Euclidean distance of each node in the two co-expression networks and the absolute number of differences of correlation coefficients between gene pairs was estimated. These three parameters were used for the network comparisons. To compare network modules, we calculated the module similarity based on the *jaccard index* among modules (Jaccard 1908).

### Diurnal variation analysis of disease modules

The 299 diseases were obtained as previously reported (Menche et al. 2015). Then the network-based distance of each gene in these diseases were calculated based on unweighted gene co-expression networks at ZT06 and ZT18. A disease module was defined based on the *P* value calculated by the 10,000 times sampling experiments in which the same number of genes were randomly selected. To detect whether the distance of disease gene sets is differential in the ZT06 and ZT18 networks, 10,000 permutation tests were performed for each disease with the same number of disease genes sampled.

### Functional enrichment analysis

The function annotation of global cycling genes was performed by g:Profiler (Raudvere et al. 2019). Each term was ranked according to the false discovery rate (FDR), and the significance threshold was set to 0.05. To assess the main function of global cycling genes, we restricted the term size to between 500 and 5000. As many enriched terms have intersections, term similarity based on the jaccard index was calculated and then clustered together by using a neighbor-joining method (Saitou and Nei 1987).

### Rhythmic interactions analysis of SARS-CoV-2 linked protein and host factors

To investigate whether rhythmic interactions exists among human SARS-CoV-2 interacting proteins, all information of 332 SARS-CoV-2 interacting proteins were downloaded as recently reported (Gordon et al. 2020). In addition, 374 SARS-Cov-2 infection host factors identified by three large-scale genome-wide CRISPR screens were collected as well. 313 interacting proteins and 333 host factors were found to be expressed in the baboon circadian transcriptome and were included in the downstream analysis. Finally, rhythmic interactions among these genes were estimated accordingly based on the baboon circadian transcriptome. The enrichment of SARS-CoV-2 interacting proteins and host factors in global cycling genes was performed using a two-sided Fisher’s exact test with odds ratio > 1 and an FDR-adjusted *P* < 0.05.

### Human interactome

The human protein–protein interactome was assembled by Zhou *et al*., (Zhou et al. 2020), which includes five types of protein–protein interactions (PPI): protein complexes data identified by AP-MS, binary protein-protein interactions tested by high-throughput yeast-two-hybrid (Y2H) systems, kinase–substrate interactions, signaling networks and literature-curated protein-protein interactions or protein 3D structures from public databases. The final dataset contains 17,706 proteins with 351,444 interactions.

### Gene sets for circadian rhythm, viral infection process and immune related functions

We downloaded viral infection process and immune related terms gene sets from AmiGO(Carbon et al. 2009). Only the human proteins are filtered. For genes that are involved in circadian rhythm, we used the “GO class (direct)” to limit them to the ones annotated directly to this function.

### Calculation of network proximity

We calculated the proximity of the SARS-CoV-2 host factors and SARS-CoV-2 interacting proteins to genes directly related to “circadian rhythm” based on previously reported methodology(Zhou et al. 2020). For the closest distance *d_AB_* between group A and group B:

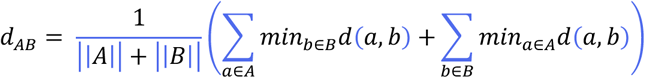

where *d*(*a*, *b*) is the shortest distance of gene a and gene b in the interactome, 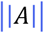 represents the size of A, 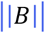 represents the size of B. To evaluate the statistical significance, 1,000 permutation tests were performed in which two randomly gene sets were chosen with the same gene number and degree distribution as A and B.

### Statistics and code availability

All the analyses were performed by R (v4.0.2). The codes for co-expression network construction are included in the supplementary document and all other codes necessary to replicate the analyses are available upon request.

## Acknowledgments

We thank John B. Hogenesch, Hiroki R. Ueda, Luoying Zhang, Hung-Chun Chang, Daniel J. Araujo and Yi Liu for helpful comments on the draft of the manuscript. This work was supported by the National Natural Science Foundation of China (Grant Nos. 31600960, 31871333, and 81827901) and the National Key R&D Program of China (Grant Nos. 2016YFC0901700 and 2016YFC1303100).

## Author contributions

Conceiving the project, G-Z.W.; performing the analyses, J.L. and P.N.; writing, reviewing and editing the manuscript, J.L., P.N., C.W.T. and G-Z.W.

